# Non-spatial features reduce the reliance on sustained spatial auditory attention

**DOI:** 10.1101/682088

**Authors:** Lia M. Bonacci, Scott Bressler, Barbara G. Shinn-Cunningham

## Abstract

Top-down spatial attention is effective at selecting a target sound from a mixture. However, non-spatial features often distinguish sources in addition to location. This study explores whether redundant non-spatial features are used to maintain selective auditory attention for a spatially defined target. We recorded electroencephalography (EEG) while subjects focused attention on one of three simultaneous melodies. In one experiment, subjects (*n* = 17) were given an auditory cue indicating both the location and pitch of the target melody. In a second experiment (*n* = 17 subjects), the cue only indicated target location, and we compared two conditions: one in which the pitch separation of competing melodies was large, and one in which this separation was small. In both experiments, responses evoked by onsets of events in sound streams were modulated equally as strong by attention, suggesting that the target stimuli were correctly selected regardless of the cue or pitch information available. In all cases, parietal alpha was lateralized following the cue, but prior to melody onset, indicating that subjects always initially focused attention in space. During the stimulus presentation, however, this lateralization weakened when pitch cues were strong, suggesting that strong pitch cues reduced reliance on sustained spatial attention. These results demonstrate that once a well-defined target stream at a known location is selected, top-down spatial attention is unnecessary to filter out a segregated competing stream.

## Introduction

Spatial features of an auditory object are often useful for focusing attention in noisy environments—if the spatial location of the object is known, then that information can be used to select this target in one location while suppressing irrelevant objects in another (Shinn-Cunningham, 2008). Often, however, additional features, such as pitch, differentiate target from distractor streams. It is therefore unclear to what extent spatial features are used when listeners must maintain attention on an auditory stream if other features also differentiate competing streams.

Selective auditory attention modulates the amplitude of event-related potentials (ERPs) in auditory cortex measured using electroencephalography (EEG); ERPs evoked by one stream are greater when that stream is attended compared to when it is ignored (Choi et al., 2013; Choi et al., 2014). Selective attention can be deployed based on a target sound’s location, or based on non-spatial features, such as pitch and timbre (Maddox and Shinn-Cunningham, 2012; Lee et al., 2012; Larson and Lee, 2014).

Spatially focused selective attention induces changes in the distribution of parietal alpha (8–14 Hz) oscillatory power. Specifically, during spatial attention, alpha power increases over parietal sensors ipsilateral to the attended location (Worden et al., 2000; Foxe and Snyder, 2011; Banerjee et al., 2011). This alpha lateralization has been studied extensively during visual spatial attention, but has been explored to a lesser degree during auditory spatial attention. As noted above, spatial attention may not be necessary to maintain attention on a target stream once it is selected based on its location. The dynamics of alpha power lateralization can thus provide insight into whether sustained attention relies on spatial processing.

In order to address this question, we measured EEG during two experiments in which subjects attended one of three competing auditory streams. Tasks were identical across experiments, but different cues were used to inform subjects as to which stream to attend. In the first experiment, an auditory cue was given that identified both the spatial location and the pitch of the target stream. Here, we asked whether subjects would orient attention in space even if they knew the pitch of the to-be-attended stream. We hypothesized that lateralization of alpha might be weak throughout attention to the cued stream since subjects did not have to orient attention in space to successfully perform the task. In the second experiment, the auditory cue only identified the spatial location of the target so that subjects would have to initially orient attention in space. We tested two conditions, presented in different blocks: one in which the pitch separation of competing melodies was large, and one in which this separation was small. We hypothesized that sustained alpha lateralization would be weak when the pitch separation was large, reflecting the fact that strong pitch cues may also be used to maintain attention to the distinct target stream, but that it would remain strong throughout trials in where spatial information was more critical for differentiating the competing streams.

## Materials and Methods

### Experimental Task and Stimuli

We conducted two separate experiments, each with the same auditory selective attention task (Fig. 1A). Three isochronous melodies were presented simultaneously from different directions—left, right, and center—using interaural time differences (ITDs) of −100 *µ*s, +100 *µ*s, and 0 *µ*s, respectively. The center melody, consisting of three 1-s notes, came on first and was always ignored. The left melody came on 0.6 s later and consisted of four 0.6-s notes. The right melody came on 0.15 s after the left melody and consisted of three 0.75-s notes. In such an arrangement, the onsets of notes in each melody were staggered in time, allowing ERPs associated with notes in each melody to be temporally isolated. In addition to being spatially separate and temporally staggered, the three melodies were separated by pitch differences, as indicated by Fig. 1B.

**Figure 1.**
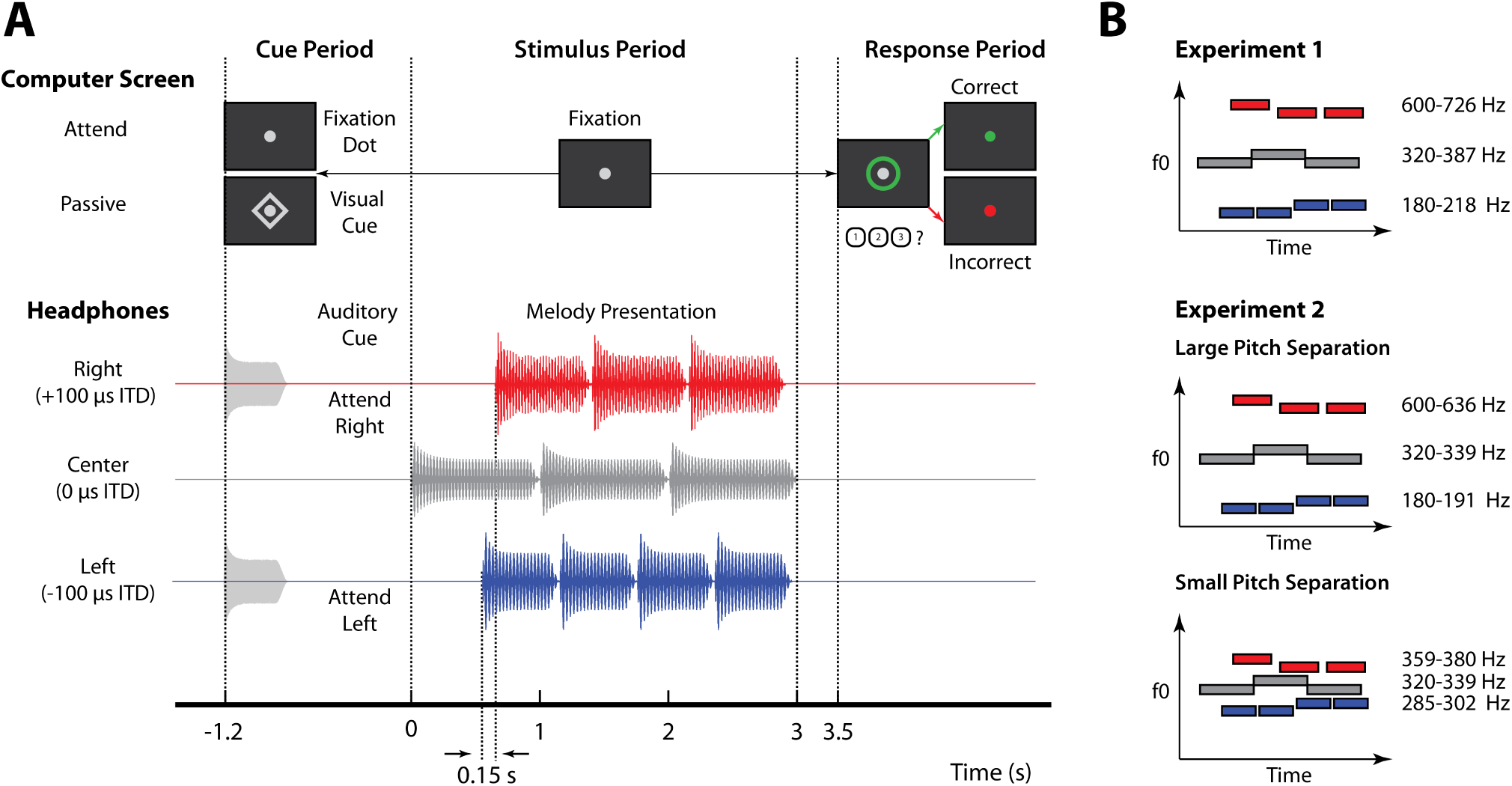
Experimental design. A, Trial structure for the auditory selective attention task. For attend-left and attend-right trials, an auditory cue was presented via headphones with the same ITD as the target melody. For passive trials, a diamond appeared around a central fixation dot on screen. During the stimulus period, subjects kept their gaze on the fixation dot while melodies were presented diotically. A green circle appeared around the fixation dot to prompt a response, and visual feedback was given after button press to indicate if the target was correctly identified. B, Left (blue), right (red), and center (grey) melodies were composed of notes with different fundamental frequencies (F0). Note that in this example, left melodies had the highest fundamentals while right melodies had the lowest fundamentals, but the opposite also occurred with equal probability. The center melody always had the same F0s, which were between F0s of the left and right melodies. Individual melodies also changed pitch over time, such that they were rising, falling, or zigzagging, illustrated by blue, red, and grey bars in this example, respectively.

Notes in each melody were composed of six harmonics added in cosine phase with magnitudes inversely proportional to frequency. Melodies were composed of two notes: a high note (H) and a low note (L). These notes were arranged to form pitch contours that were “rising”, “falling”, or “zigzagging”. “Rising” melodies started on the low note and transitioned at a randomly-selected point to the high note (e.g., L-L-H-H). “Falling” melodies started on the high note and transitioned at some onset to the low note (e.g., H-L-L-L). “Zigzagging” melodies started on either the high or low note, transitioned to the opposite note, and then returned to the starting note (e.g., L-H-L or H-L-H). In “zigzagging” melodies, the second pitch change always occurred between the last two notes to ensure subjects had to maintain focused attention for the duration of the auditory stream. Contours were selected independently for left, right, and center melodies, with each contour having a 1/3 chance of being chosen.

At the beginning of each trial, subjects were given an auditory cue directing them to attend either the left or the right melody. After attending the target melody, subjects had to report its pitch contour via button press. In addition to active attention trials, passive trials were included in which subjects were given a visual cue, signaling they could ignore stimuli and were to withhold a response. All cues were 100% valid. Visual feedback was given at the end of each trial to indicate if the melody was correctly identified.

Subjects performed the experiment in front of an LCD monitor in a sound-treated booth. Stimuli were generated using MATLAB (MathWorks, Natick, MA) with the PsychToolbox 3 extension (Brainard, 1997). Sound stimuli were presented diotically via Etymotic ER-1 insert headphones (Etymotic, Elk Grove Village, IL) connected to Tucker-Davis Technologies System 3 (TDT, Alachua, FL) hardware which interfaced with MATLAB software that controlled the experiment. During the task, subjects were instructed to keep their eyes open and to foveate on a central fixation dot.

#### Experiment 1

In Experiment 1, the auditory cue was a six-harmonic complex tone that was presented with the same ITD as the target melody. The fundamental frequency of this cue was also in the same pitch range as the notes composing the target. As mentioned above, each melody was presented in a different pitch range, as shown in Fig. 1B. Within each pitch range, two of three possible fundamental frequencies were randomly selected to compose the high and low note for each two-note melody. The three possible fundamentals were separated by 1.65 semitones. The center melody, which was always ignored, had notes with fundamentals in the 320–387 Hz range. On a given trial, either the right or left melody was selected, with equal probability, to have fundamentals in the 180–218 Hz range. The remaining melody was selected to have fundamentals in the 600–726 Hz range. This structure ensured that each melody was perceptually segregated from the others.

Trials were arranged in 9 blocks of 30, with each block containing 1/3 attend-left and 1/3 attend-right trials presented in random order. The remaining trials were passive control trials. This resulted in 90 trials for each condition. Before performing the task, subjects were required to pass a training demo in which they were presented with a series of single melodies and asked to identify their pitch contours. Passive trials were also included in the training demo to ensure subjects knew when to withhold a response. In order to continue the study, subjects had to answer correctly on 10 of 12 demo trials (4 passive trials, 8 active attention trials). This requirement was included to ensure that subjects’ performance on the task was not limited by their ability to identify pitch contours, but by their ability to direct attention.

#### Experiment 2

In experiment 2, the auditory cue was a white noise burst that was presented with the same ITD as the target melody. This required subjects to at least initially orient attention in space since no pitch information was available in the cue. As in Experiment 1, each melody was presented in a different pitch range. Within each pitch range, the same two fundamentals were used to compose the high and low note of each two-note melody. These two fundamentals were separated by 1 semitone. In all trials, the center melody always had fundamentals in the middle, 320–339 Hz range. As in experiment 1, high and low pitch ranges were randomly assigned to the left and right melodies.

The fundamental frequency of melodies in these pitch ranges depended on the experimental block, which were one of two conditions: one in which the pitch separation of competing melodies was large and one in which it was small (Fig. 1B). In the large pitch separation condition, the low pitch melodies had fundamentals in the 180–191 Hz range while the high pitch melodies had fundamentals in the 600–636 Hz range, creating clearly segregated streams. In the small pitch separation condition, fundamentals of low (285–302 Hz) and high (359–380 Hz) pitch ranges were shifted closer to that of the center melody. The resulting sound mixture was thus more difficult to automatically segregate by pitch alone. Large and small pitch separation blocks were grouped together in pairs, but the order of conditions was random for each pair of blocks (e.g., Lg-Sm-Sm-Lg-Sm-Lg-Lg-Sm).

Trials were arranged in 16 blocks of 30, with each block containing 2/5 attend-left, 2/5 attend-right, and 1/5 passive trials. This resulted in 96 attend-left and attend-right trials in each pitch separation condition, and 96 passive trials across all pitch separation conditions. After the first 8 blocks, subjects were instructed to take a break before starting the remaining set of 8 blocks. As in Experiment 1, subjects were required to pass a training demo in which they had to identify the pitch contour of a single melody presented alone. Two training blocks were given, one each for stimuli in the two pitch separation conditions. Each block contained 15 trials (3 passive trials, 12 active attention trials), and subjects had to answer correctly on 13 trials for each block to continue in the experiment.

### Subjects

Data from a total of 34 subjects with normal hearing and no known neurological disorders were analyzed as part of this study—17 for Experiment 1 (8 male, mean age=21.88, *SD* = 2.78) and 17 different subjects for Experiment 2 (9 male, mean age=22.35, *SD* = 3.67). Additional subjects performed both experiments (four subjects for Experiment 1, and three subjects for Experiment 2), but produced data that had to be discarded due to too many incorrect-response trials or too many trials with noisy EEG. An audiogram was conducted for each subject to confirm that thresholds were below 20 dB HL at octave frequencies from 250 Hz to 8 kHz. Some subjects recruited for Experiment 2 were dismissed early from the study: one had audiometric thresholds above the required level, two could not give a clean EEG signal, and six failed the training demo described above. These subjects were compensated for their time, but did not have EEG recorded. All subjects gave written informed consent prior to participation and were compensated at an hourly rate ($25/hr for Experiment 1, $15/hr for Experiment 2) as well as with a bonus for each correct response ($0.02 per response, up to $7.50 per hour). All procedures were approved by the Boston University Institutional Review Board.

Subjects who participated in Experiment 1 also participated in an analogous visual task—not described here—during the same experimental session. Of these subjects, 12 participated in the visual task after the auditory task was complete. The remaining 5 subjects completed the visual task blocks first. Subjects who participated in Experiment 2 were not exposed to any visual analog of the task. While it is possible that the subjects who completed visual experiment before starting the auditory experiment were biased towards using spatial features for selection, we found no evidence that alpha modulation was statistically different between the subjects who completed the visual task first and those who did not.

### Data Collection

EEG data were recorded at a sampling rate of 2048 Hz using the BioSemi ActiveTwo system and its ActiveView acquisition software (BioSemi, Amsterdam, Netherlands). A 64-channel cap with electrode positions arranged according to the international 10-20 system was used for measurement, along with two reference electrodes placed on the mastoids. An additional three electrodes were placed around the eyes for electrooculogram (EOG) measurement, which was used to detect eye blinks for later removal from the EEG signal. Event triggers were driven by MATLAB software running the experimental task and generated by Tucker-Davis Technologies System 3 (TDT, Alachua, FL) hardware that interfaced with the computer recording EEG data. In Experiment 2, RME Fireface UXC hardware was used instead of the TDT for trigger generation. An EyeLink Plus 1000 (SR Research, Ottawa, Ontario, Canada) eye tracker was used in Experiment 2 to ensure subjects did not close or move their eyes during the task. In Experiment 1, subjects were instructed to fixate on a central fixation dot, but eye tracking was not recorded or monitored during the experiment.

### Data Analysis

#### EEG Processing

The EEGLAB toolbox for MATLAB (Delorme and Makeig, 2004) was used to process raw EEG data. Raw EEG data were first re-referenced to the average between two mastoid electrodes and downsampled to 256 Hz. An FIR zero-phase filter with cutoffs at 1 and 20 Hz was then applied to the signal. Eye blinks were removed using independent component analysis. Epochs with amplitudes over ±100 *µ*V were rejected along with trials in which subjects gave an incorrect response. CSD Toolbox (Kayser and Tenke, 2006) was used to transform EEG data from voltage to current source density. This technique was employed to reduce spatially correlated EEG noise, which is desirable when localizing parietal alpha power across the scalp (McFarland, 2015; Kayser and Tenke, 2015).

#### Event-Related Potential

The ERP time course was estimated using a bootstrap procedure. The average time course was first calculated across 100 randomly chosen trials with replacement within a single subject and condition. This procedure was repeated 200 times, and each subject’s estimated ERP was taken as the average across these 200 trials. Each subject’s ERPs were then normalized by dividing the entire time series by the average amplitude of the N1 response to the first distractor onset, averaged across all trials and channels. This step ensured that all ERPs were similar in magnitude across subjects. The first distractor onset was selected since it was previously shown to elicit a strong N1 response that is not modulated by attention (Choi et al., 2014), presumably due to the salience of the initial sound onset eliciting involuntary attention. Grand averages were obtained for each condition by averaging the normalized ERP amplitudes across subjects.

N1 amplitudes were extracted for each subject from the normalized ERP time courses. These ERPs were first averaged across 17 frontocentral channels where responses were largest (Fp1, Fp2, AF3, AF7, AF4, AF8, F1, F3, F5, F7, F2, F4, F6, F8, Fpz, AFz, Fz). This normalized channel average was then averaged across subjects in order to estimate the N1 timings for each note onset. These times, selected based on the largest negative value of the ERP in a window between 75 and 240 milliseconds following each stimulus onset, were then used to estimate N1 amplitudes for each subject’s channel-averaged ERP. The ERP was averaged in a 50-ms window centered around each of the selected time points to quantify N1 amplitude in response to each note. Each subject’s ERP was visually inspected to ensure that N1s were correctly identified.

Attentional modulation of the N1 was quantified for each subject using an attentional modulation index, AMI_N1_, given by Eq. 1.

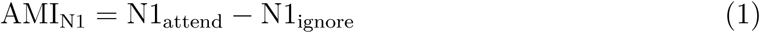

Here, N1_attend_ is the N1 amplitude elicited by the onset of a particular note when it was attended; N1_ignore_ is the N1 amplitude elicited by the same note when it was ignored. AMI_N1_ was calculated for each note in both left and right melodies and averaged to quantify overall modulation of the N1. The N1 to the first leading left onset was not included in this average since since it elicited a strong automatic response regardless of cue condition, as in (Choi et al., 2014). Large positive values of AMI_N1_ indicate that N1s were overall larger when notes were attended, relative to when they were ignored.

#### Induced Alpha Power

To obtain the induced alpha response, it was necessary to first remove phase-locked activity. To achieve this, the phase-locked, or evoked response (ERP) was calculated as described above and subtracted from each trial, leaving only the induced, non-phase-locked activity. Power at each frequency in the alpha band (8–14 Hz) was estimated for each trial using a short-time Fourier transform. An individual alpha frequency (IAF) was selected for each subject by finding the frequency in the range of 8–14 Hz with the greatest magnitude across attend-left and attend-right conditions in 20 parietal and occipital channels (P2, P4, P6, P8, P10, PO4, PO8, O2, P1, P3, P5, P7, P9, PO3, PO7, O1, Pz, POz, Oz, Iz). Power was extracted at this IAF to produce a single time series for each trial in each EEG channel. The bootstrap procedure described above for estimating the ERP was used to estimate each subject’s average induced alpha power for each experimental condition. Normalization was performed on individual subject trial-averaged time series by dividing each time point by that subject’s average alpha power across time, sensors, and experimental condition. Grand averages were obtained by averaging these normalized time series across subjects. Quantities shown on topoplots represent averages across the cue period (−1.2–0 s) or stimulus period (0–3 s).

An attentional modulation index of alpha power, AMI_*α*_, was also calculated for each subject. Calculation of AMI_*α*_ is given by Eq. 2.

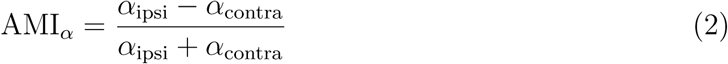

In Eq. 2, *α*_ipsi_ is the average alpha power during the stimulus or cue period, measured ipsilateral to the cued sequence, or rather contralateral to the ignored sequence; *α*_contra_ is this average alpha power, measured contralateral to the cued sequence. Large positive values of AMI_*α*_ indicate that alpha power was overall larger ipsilateral to cued stimuli (i.e., the alpha response was larger over cortices processing ignored information). Averages were calculated across left and right parietal and occipital channels separately, depending on the attention condition (i.e., left channels for *α*_ipsi_ in attend-left trials and right channels for *α*_ipsi_ in attend-right trials). These averages were then collapsed across attention conditions and parietal sensors to quantify *α*_ipsi_ and *α*_contra_.

#### Significance Testing

For Experiment 1, we performed statistical testing to determine if modulation of N1 was significantly greater than zero. For this purpose, we used a one-sample, one-sided *t*-test on AMI_N1_ data. We also wanted to determine if alpha lateralization, indexed by AMI_*α*_, was significantly greater than zero in both the cue and stimulus periods. Again we used a one-sample, one-sided *t*-test. In order to correct for multiple comparisons, Bonferroni and Benjamini-Hochberg criteria were used to determine significance at the *α* = 0.05 significance level. The same statistical procedures were used in Experiment 2 to determine if AMI_N1_ and AMI_*α*_ were significantly greater than zero. We also hypothesized that AMI_*α*_ would be greater in the small pitch separation condition than in the large pitch separation condition. To determine if this difference was significant, we performed paired-sample, one-sided *t*-tests for AMI_*α*_ values measured during cue and stimulus periods. Multiple comparisons procedures were performed before determining significance. AMI_N1_ was also compared between large and small pitch separation conditions using a paired-sample *t*-test.

## Results

### Behavior

#### Differences in performance existed between attend-left and attend-right trials in Experiment 2

Performance, measured as percent correct response, is displayed for both experiments in Fig. 2. Overall, subjects performed well above chance, suggesting successful focus of attention. In Experiment 1, no significant differences were found between attend-left and attend-right trials (*p* = 0.24, paired *t*-test). Differences in performance were found in Experiment 2, however. In the large pitch separation condition, subjects performed better on attend-right trials than attend-left trials (*p* = 0.005, paired *t*-test). The opposite was true for the small pitch separation condition (*p* = 0.013, paired *t*-test). In comparing spatial attention conditions across pitch separation conditions, there was no significant difference in performance on attend-left trials between large and small pitch separation conditions (*p* = 0.94, paired *t*-test). For attend-right trials, however, performance was significantly greater in the large pitch separation condition (*p* < 0.001, paired *t*-test).

**Figure 2.**
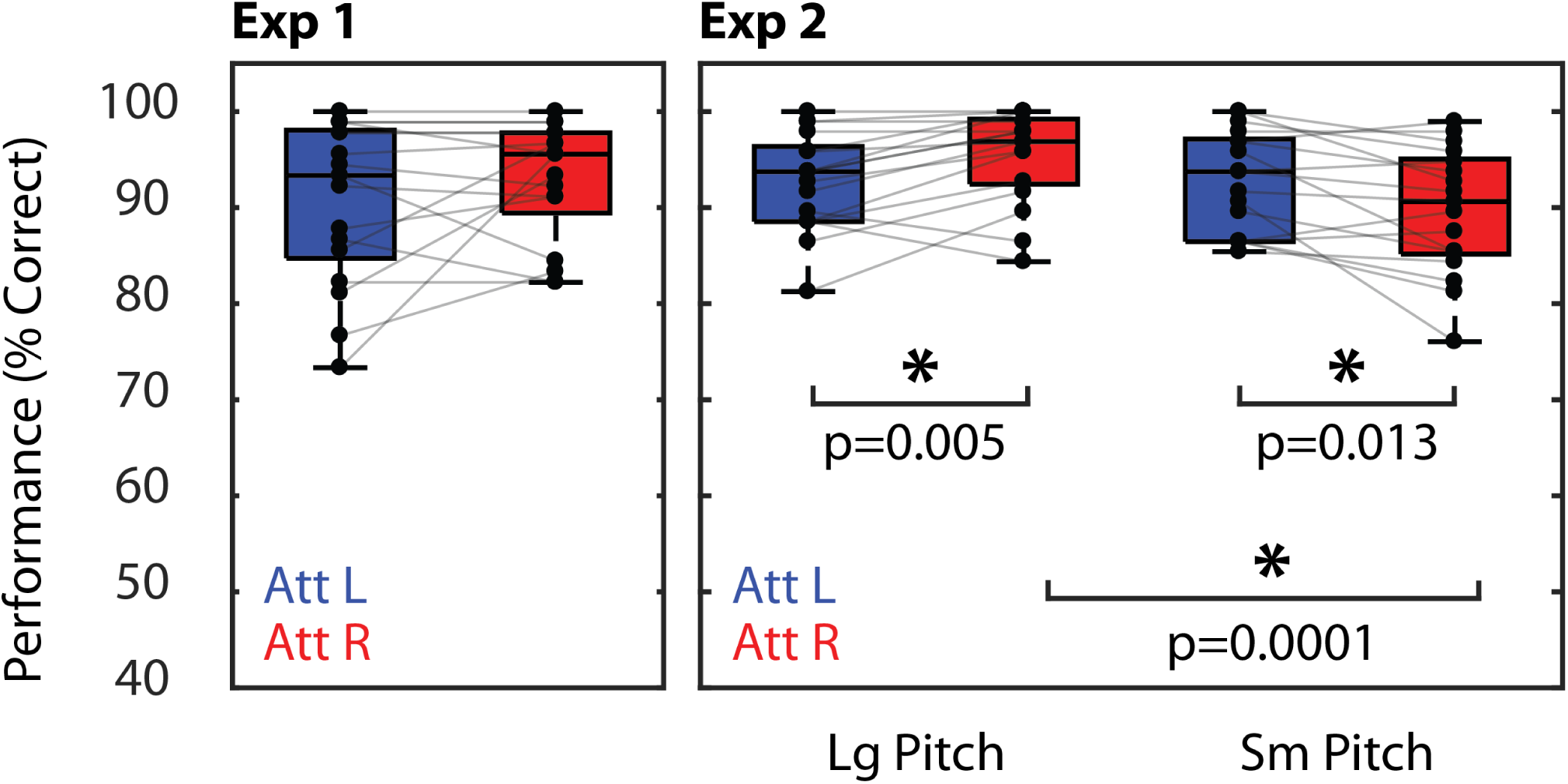
Percent correct scores for Experiments 1 (left) and 2 (right). Asterisks indicate significant differences between conditions (*p* < 0.05, *t*-test). No corrections were made for multiple comparisons.

These performance differences may be explained by differences in the bottom-up salience of the melodies in the two conditions. Recall that the right melody always lagged the left melody in time. Therefore, in the large pitch separation condition, even though the leading (left) melody may have captured attention first, the right melody had a distinctive pitch that caused the lagging melody to be heard as a new event automatically. In the small pitch separation condition, the lagging melody had a similar pitch to the leading melody, which likely made the melody onset less clear and salient.

### Event-Related Potential

#### In both experiments, the N1 response was similarly modulated by selective attention

In Experiment 1, N1 amplitudes were modulated by attention (Fig. 3, top panel). When subjects were cued to attend the left melody, N1 amplitudes were more negative in response to left note onsets (blue vertical lines) when those notes were attended (blue trace) than when they were ignored (attend-right trials, red trace). Similarly, when subjects were cued to attend the right melody, the N1 was more negative at right note onsets (red vertical lines) when those notes were attended than when they were ignored. The same modulation of the N1 was observed in Experiment 2, both in the large pitch separation condition (middle panel) and the small pitch separation condition (bottom panel). This modulation was quantified using the attentional modulation index, AMI_N1_ described above. In both experiments, AMI_N1_ was significantly greater than zero (*p* < 0.001, *t*-test), indicating that the N1s from auditory cortex were always larger in response to attended stimuli than ignored stimuli. AMI_N1_ was also compared between pitch separation conditions in Experiment 2, but no significant difference in modulation was found (*p* = 0.26, *t*-test). Thus, the degree of N1 modulation did not change significantly based on the degree of pitch information available in this experiment, suggesting that subjects selected target stimuli regardless of the available pitch cues.

**Figure 3.**
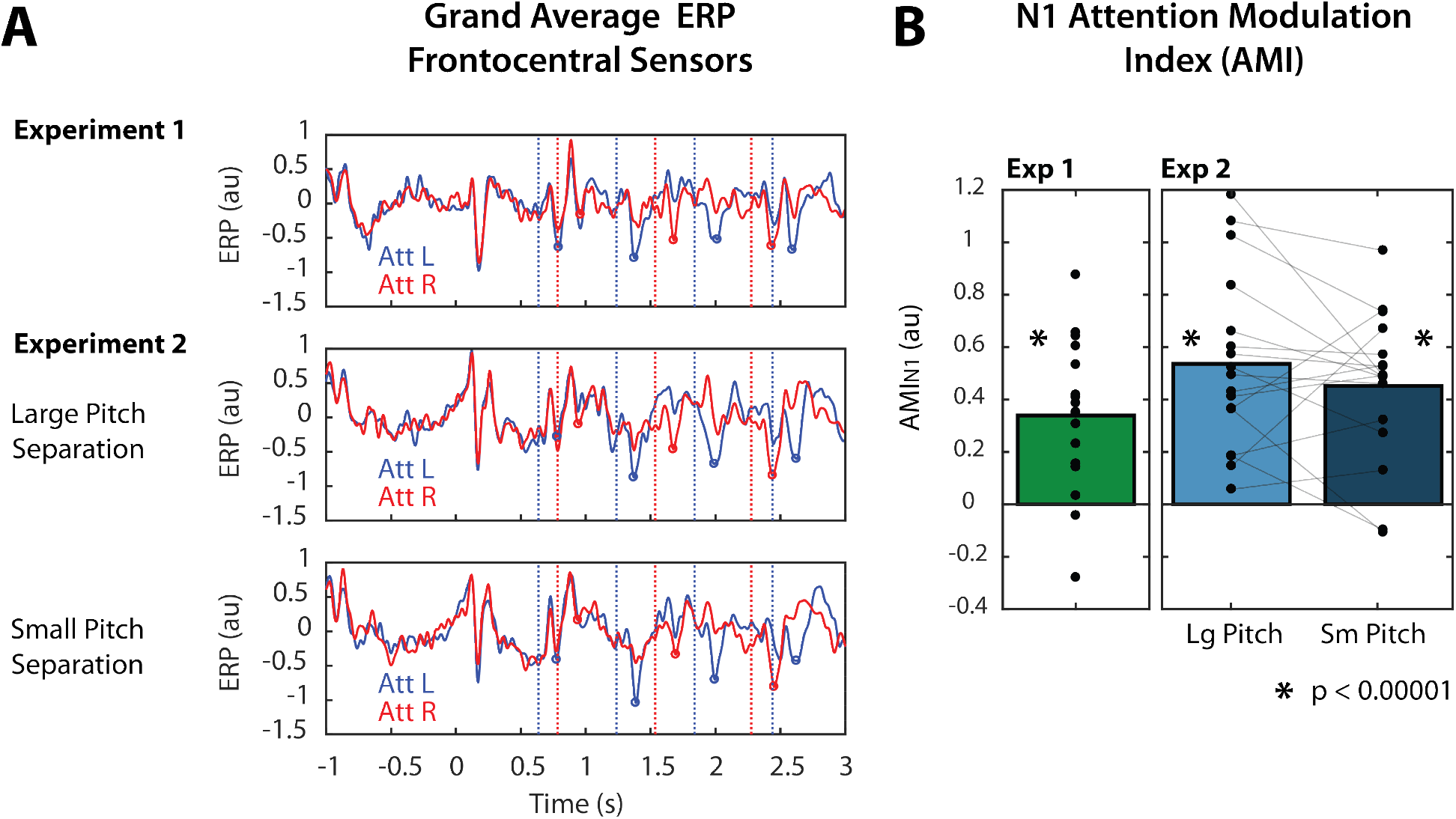
A, Grand average (n=17) normalized ERP responses over time in Experiment 1 (top) and Experiment 2 (bottom). ERPs were averaged across frontocentral EEG sensors. Red and blue vertical lines indicate right and left note onset times, respectively. Red and blue circles indicate the identified N1 peak amplitudes in response to right and left notes, respectively. B, N1 modulation summarized as AMI_N1_. Individual points indicate individual subject AMI_N1_. Asterisks indicate that AMI_N1_ was significantly greater than zero at the *p* = 0.0001 significance level (one-sided, one-sample *t*-test). No corrections were made for multiple comparisons.

### Induced Alpha Power

#### During the cue period, alpha power was always lateralized across parietal sensors

Grand average alpha power differences, averaged over the cue period, are shown in the left panels of Fig. 4A and Fig. 4B for both experiments. Figure 4A shows alpha power differences between attend-left and attend-right trials. In all cases, average alpha power was greater in left parietal sensors during attend-left trials than during attend-right trials. Similarly, in right parietal sensors, alpha power was greater in attend-right trials than in attend-left trials. Figure 4B shows these differences collapsed across left and right parietal sensors, so that alpha is represented as the difference between ipsilateral and contralateral attention conditions. During the cue period, alpha power was greater when attended stimuli were ipsilateral to a given parietal sensor than when attended stimuli were contralateral to that same sensor. This suggests that alpha increased contralateral to the ignored location, supporting the idea that alpha reflects suppression of distractors.

**Figure 4.**
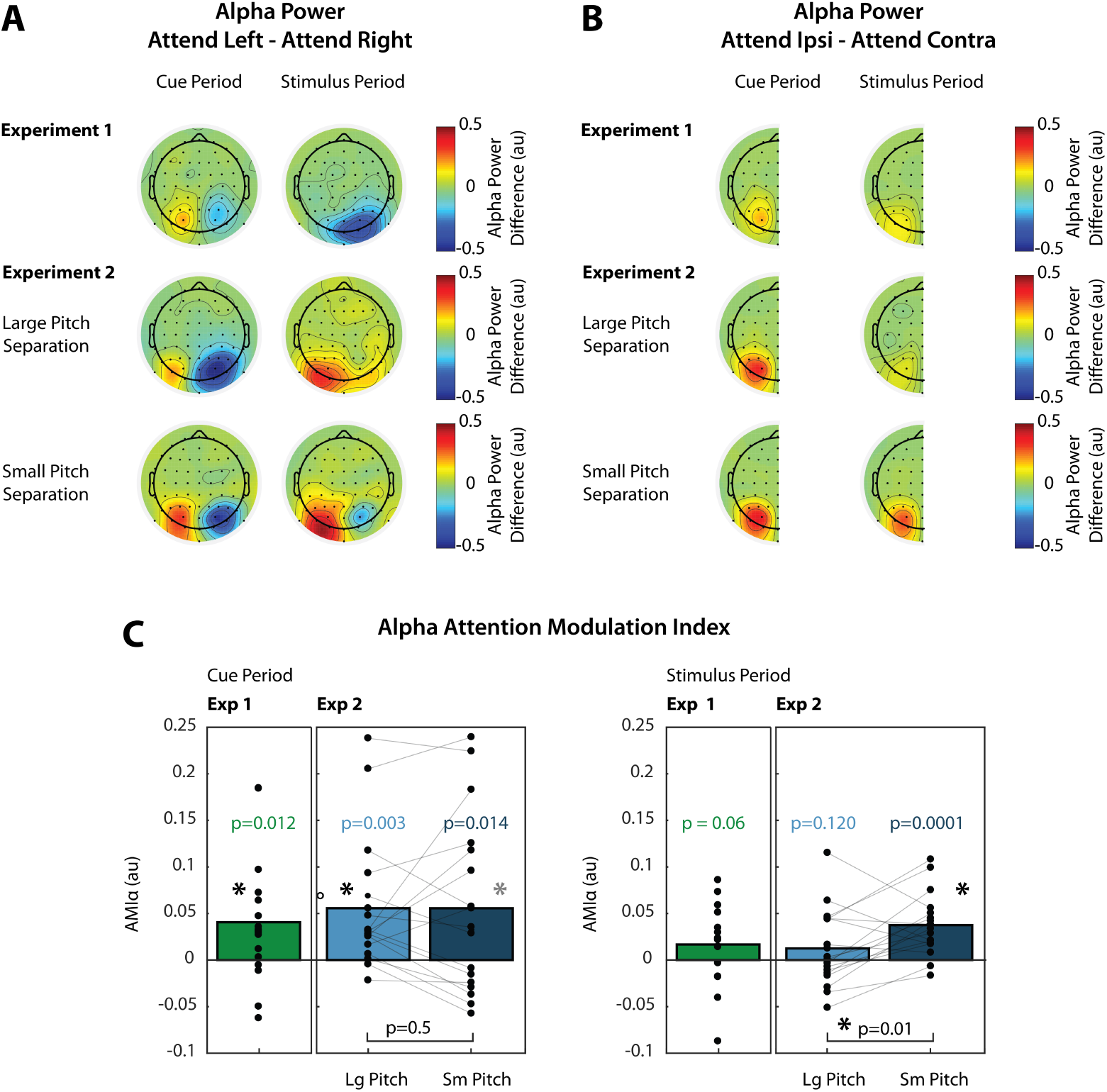
A, Grand average (n=17) normalized alpha power differences between attend-left and attend-right trials. For each channel, alpha power was averaged across time during the cue period (left, t = −1.2–0 s) or during the stimulus period (right, t = 0–3 s). B, Grand average alpha power differences between ipsilateral and contralateral attention conditions, collapsed across left and right parietal channels. C, AMI_*α*_ calculated during the cue period (left panel) and stimulus period (right panel). Displayed *p*-values were not corrected for multiple comparisons. Asterisks over individual bars indicate that AMI_*α*_ was significantly greater than zero after correcting for multiple comparisons using Bonferroni (black asterisks) or Benjamini-Hochberg (grey asterisks) procedures. Comparisons between conditions in Experiment 2 are also shown by brackets and associated *p*-values. Again, asterisks indicate significant differences after correction for multiple comparisons.

Figure 4C shows attention modulation indices (AMI_*α*_), which are based on the ipsilateral/contralateral differences shown in Fig. 4B. In Experiment 1, alpha was lateralized during the cue period (green bars), and this lateralization was significantly greater than zero (*p* = 0.012, *t*-test). In experiment 2, AMI_*α*_ was also measured during the cue period, and was significantly greater than zero for both the large (light blue bars) and small (dark blue bars) pitch separation conditions (*p* = 0.003 and *p* = 0.014, respectively, *t*-test). When the Benjamini-Hochberg procedure was applied to control the false discovery rate (FDR=0.05) in each experiment, these values of AMI_*α*_ remained significantly greater than zero. When Bonferroni correction was applied, AMI_*α*_ during the cue period was still significantly greater than zero in Experiment 1 (*p* < 0.025) and in the large pitch separation condition in Experiment 2 (*p* < 0.0125). In experiment 2, there was no significant difference in AMI_*α*_ between large and small pitch separation conditions (*p* = 0.5, paired *t*-test). These results suggest that subjects always initially oriented attention using known spatial features of the target.

#### During the stimulus period, alpha lateralization was weak when pitch cues were strong

Grand average alpha power differences, averaged over the stimulus period, are shown in the right panels of Fig. 4A and Fig. 4B for both experiments. While alpha power was always lateralized during the cue period, this lateralization only persisted during the stimulus period in the small pitch separation condition of Experiment 2. Here, alpha power was larger in left parietal sensors during attend-left trials and larger in right parietal sensors during attend-right trials. In the large pitch separation condition, alpha power was larger in left parietal sensors during attend-left trials. In right parietal sensors, alpha was also greater during attend-left trials, but this difference was smaller than in left parietal sensors. In Experiment 1, alpha power in right parietal sensors was greater during attend-right trials. In left parietal sensors, there was not a large difference between attend-left and attend-right trials.

Figure 4B shows these differences collapsed across parietal sensors. Here, we see that there was not an overall difference in alpha lateralization between ipsilateral and contralateral attention trials during the stimulus period in Experiment 1 or in the large pitch separation condition of Experiment 2. In the small pitch separation condition, the difference between alpha power in ipsilateral and contralateral attention trials was similar to that observed during the cue period. These differences are represented as AMI_*α*_ in Fig. 4C. While AMI_*α*_ was not significantly greater than zero in Experiment 1 (*p* = 0.06, *t*-test) or in the large pitch separation condition (*p* = 0.12, *t*-test), it was significantly greater than zero in the small pitch separation condition (*p* < 0.001, *t*-test). When Bonferroni criteria were applied to correct for multiple comparisons, AMI_*α*_ was still significantly greater than zero in the small pitch separation condition of Experiment 2 (*p* < 0.0125). We also determined that AMI_*α*_ was significantly larger in the small pitch separation condition compared to the large pitch separation condition (*p* = 0.01, paired *t*-test), and this difference was still significant after correcting for multiple comparisons (*p* < 0.025, Bonferroni).

#### AMI_*α*_ was not correlated with performance measures

We asked whether the differences in alpha modulation between large and small pitch separation conditions could be explained by the performance differences described above. We therefore looked for correlations between AMI_*α*_ measures and percent correct scores. For alpha power, we calculated AMI_*α*_ separately for left and right parietal channels and looked for correlations with percent correct scores in attend-left or attend-right trials (4 comparisons in each pitch separation condition). We found no significant correlation between any combination of AMI_*α*_ and percent correct scores (Spearman rank correlation, *p* > 0.2 for all comparisons).

## Discussion

### Modulation of the N1 response reflects selection of target stimuli, but does not suggest which feature was used to perform selection

In both experiments, we observed similar modulation of the N1 response in frontocentral channels, suggesting that there was no effect of pitch cue strength on N1 modulation. Modulation of N1s from auditory cortex reflects enhancement of target stimuli as well as suppression of distracting stimuli (Choi et al., 2014; Kong et al., 2014). Therefore, the fact that we observed no difference in N1 modulation between experimental conditions suggests that subjects were able to focus attention on the target stream, even when pitch differences were small. A previous study that used a similar paradigm (Choi et al., 2014) found that when competing melodies were in overlapping pitch ranges, N1 modulation was degraded and performance was significantly worse than when melodies were in separate ranges, exactly as in Experiment 1 here. This was likely due to difficulty segregating the competing streams. The fact that we did not observe degraded N1 modulation in the small pitch separation condition was likely due to the fact that competing melodies did not have overlapping pitch ranges as in (Choi et al., 2014), but distinct ranges that were close together (∼1 semitone difference). This design difference, and the fact that behavioral measures show that subjects had performed well on the task in all conditions, suggests that subjects were able to segregate and select targets regardless of the available pitch cues.

The fact that the N1 was modulated similarly does not mean that spatial features were used in the same way to maintain attention across conditions. In fact, there are a number of experiments that show N1 modulation in response to attended auditory stimuli that were not spatially separate from competing objects (Hansen and Hillyard, 1988; Kong et al., 2014). Thus, the N1 serves as an index of selective attention independent of the features used for selection. If we wish to index the extent to which spatial features are used to direct top-down attention, then measuring the N1 is insufficient if other features can also be used. Instead, we look to modulation of alpha power, which occurs over cortical regions that map space. If spatial features are used to a lesser degree to focus attention, then we may expect reduced attentional modulation of parietal alpha power.

### Lateralization of parietal alpha power reflects spatial focus of auditory selective attention

While parietal alpha has been studied extensively as a correlate of visuospatial attention, its role in auditory spatial attention is less clear. Nonetheless, growing evidence supports the idea that auditory spatial attention recruits the same cortical networks that are active during visual spatial attention. Early neuroimaging studies defined a dorsal frontoparietal network responsible for orienting visual attention to a particular location (Posner and Petersen, 1990; Corbetta and Shulman, 2002; Petersen and Posner, 2012). This network was composed of the frontal eye fields (FEF) and superior parietal lobe (SPL). Later studies revealed that this network is also involved in auditory attention (Lewis et al., 2000; Shomstein and Yantis, 2006; Krumbholz et al., 2009; Braga et al., 2013), but did not establish whether the network was truly supramodal or was instead composed of modality-specific subnetworks.

Recent fMRI studies have identified interleaved visual and auditory-biased networks in lateral frontal cortex (LFC) (Michalka et al., 2015; Noyce et al., 2017), suggesting that there are modality-specific networks for attention. The visual-biased network contains superior and inferior precentral sulcus (iPCS and sPCS), which are functionally connected to posterior visual sensory regions; the auditory-biased regions contain transverse gyrus intersecting precentral sulcus (tgPCS) and caudal inferior frontal sulcus (cIFS), which are functionally connected to posterior auditory sensory regions. Although these networks are modality-specific, the visual-biased network is flexibly recruited during auditory attention when spatial focus is required to perform the task (Michalka et al., 2015; Michalka et al., 2016; Noyce et al., 2017). When the task has high temporal demands, the auditory-biased network is active in both vision and audition. Thus, while there are modality-specific networks for attention, these networks are recruited in a non-modality specific manner depending on the attended features (spatial vs. temporal).

If the same frontoparietal network underlies auditory and visual spatial attention, then should expect to observe the same EEG correlates of spatial attention over parietal cortex during spatial attention independent of stimulus modality. Therefore, if increased parietal alpha reflects suppression of unattended space in vision (Worden et al., 2000; Sauseng et al., 2005; Kelly et al., 2006; Foxe and Snyder, 2011; Händel et al., 2011; Payne et al., 2013), then it should be present for spatial suppression in audition. Indeed, at least one previous study has shown evidence of parietal alpha modulation during auditory spatial attention (Banerjee et al., 2011).

Our results are consistent with these findings. We observed that alpha was lateralized after subjects were given a spatial cue, and this lateralization pattern reflected the space being ignored (i.e., alpha was greater ipsilateral to the attended location). The results from Experiment 1 suggest that subjects at least initially oriented top-down attention using known spatial features of the target even if they could depend solely on pitch information to perform the task. In Experiment 2, subjects *had* to initially orient attention in space due to the absence of pitch cues. Therefore, the observed alpha modulation during the cue period in this experiment strengthens the argument that parietal alpha lateralization reflects the use of *spatial* features to help focus attention.

### Alpha lateralization is weak when pitch cues are strong, reflecting the fact that pitch can also be used to help focus attention

While space is first coded at the level of the retina in vision, the auditory system relies on interaural time and level differences to localize sound. Therefore, the mechanisms by which auditory attention operate are likely not inherently spatial. This explains why in the vision literature, spatial and feature-based (e.g., color, texture, etc.) attention are described separately, yet in audition, perceived location is described as a feature itself (Shinn-Cunningham, 2008; Maddox and Shinn-Cunningham, 2012; Shinn-Cunningham et al., 2017). In audition, non-spatial features, such as pitch, can often be used to direct and maintain attention to an ongoing stream (Maddox and Shinn-Cunningham, 2012; Lee et al., 2012; Lee et al., 2014). If these pitch cues are large compared to the available spatial cues, then individuals may depend more on pitch as the feature on which to base attention. When pitch cues are less informative, however, it may be more beneficial to depend on spatial differences among competing stimuli to maintain attention.

If parietal alpha truly reflects the use of spatial features during sustained top-down attention, then its modulation should be weaker during tasks in which spatial features are redundant with other non-spatial features. Our results support this view. As argued above, alpha lateralization occurred during the cue period in all conditions, which suggests that spatial attention was initially directed using the known spatial features. During the stimulus period, however, this lateralization was weak (i.e., not significantly greater than zero) when strong pitch cues were available (i.e., Experiment 1 and the large pitch separation condition of Experiment 2). In Experiment 2, we also observed that this lateralization was significantly larger in the small pitch separation condition than in the large pitch separation conditions. These results likely reflect the fact that, in addition to space, pitch cues could also be used to differentiate target from distractor. Therefore, even though subjects initially directed attention to the location of interest, once the auditory object was selected, its pitch was used to maintain attention throughout the remainder of the stream. When these pitch cues were weak, spatial features may have been necessary to maintain attention, which is why we observed alpha lateralization throughout the small pitch separation trials.

### The degree of alpha lateralization doesn’t explain performance

In Experiment 2, we observed differences in performance between pitch separation conditions. Therefore, it may be possible that the differences in alpha lateralization observed between the two conditions are due to differences in ability to perform the task instead of differences in pitch cue strength. However, we argue this is not the case for two reasons. First, we removed all trials in which subjects responded incorrectly, so we assume the EEG signal we observed was recorded when subjects successfully focused attention and not when they may have been struggling to do so. Second, if it *were* the case that alpha lateralization was stronger because the small pitch separation condition was more difficult, then we may expect some correlation of AMI_*α*_ with performance measures—subjects who are inherently worse at the task may require more suppression of distractors, which may manifest in greater alpha lateralization. However, we observed no correlation of performance measures with alpha modulation in either left or right parietal channels. Furthermore, that a similar lateralization pattern was observed during the cue period for both large and small pitch separation conditions despite the performance differences suggests that alpha is indexing spatial focus of attention and not task difficulty.

One may still argue that the greater alpha lateralization observed during the stimulus period was due to more effort being required in the small pitch separation condition, even though performance measures were not correlated with lateralization measures. While this may be true, we argue here that the increased effort required may be defined as the greater need to orient and maintain attention in space since pitch cues are less informative. Therefore, more effort here means more use of spatial attention, which is reflected by stronger alpha lateralization. In the future, more efforts should be made to disentangle the effects of task difficulty and spatial attention on parietal alpha power.

### Caveats: Weighting the Effects of Pitch and Space

While our results suggest that the degree of alpha lateralization reflects the degree to which spatial features are used to selectively attend, we did not parametrically adjust spatial and pitch separations of competing melodies. Rather, we tested two conditions in which the pitch separation was different while a somewhat small spatial separation (±100 *µ*s ITD) was held constant. Therefore, in this study, we assumed that when pitch differences are larger, less dependence on spatial features is required during the course of the auditory stream. It may be the case, however, that given the same pitch separation, spatial features would be used to a greater extent if the spatial separation were larger. In fact, previous studies have shown that both pitch and perceived location have similar effects on ability to selectively attend (Maddox and Shinn-Cunningham, 2012), with performance improving as the task-relevant feature separation increased. Future work should aim to address under what conditions and to what degree alpha is lateralized given the available space and pitch cues.

## Conclusions

Understanding which features are used during auditory attention advances our understanding of how individuals communicate in noisy environments. In these settings, individuals must not only be able to segregate various auditory objects, but also select a single relevant object while suppressing distractors. Our results show that while N1 modulation reflects selection during both spatial and non-spatial auditory attention, parietal alpha lateralization reflects the degree to which spatial features are used to suppress distractors. This EEG correlate that reflects the degree to which spatial features are used could be a candidate feature when designing a non-invasive method for tracking the use of spatial features in a variety of complex auditory scenes—scenes which contain different levels of spatial and non-spatial features. In addition to behavioral studies, such EEG studies could reveal under which conditions spatial features are used to help solve the cocktail party problem (Cherry, 1953).

Communication at the cocktail party is particularly challenging for those who are hearing impaired, even when assistive devices are worn (Marrone et al., 2008). There is a need for technologies that assist object selection in complex scenes. Proposed strategies involve predicting where individuals intend to direct attention in order to enhance selection of objects at that location (Shinn-Cunningham and Best, 2008; Kidd Jr et al., 2013). Such predictions may be made using measures from non-invasive EEG. However, in order to predict *where* individuals intend to focus attention using correlates of spatial attention, we have to know that spatial attention is being used in the first place. Our results suggests that the degree of parietal alpha lateralization may reflect the degree to which spatial features are used during attention, and so if an individual is attempting to orient attention using non-spatial features, then alpha lateralization would not be informative. This technique would instead have to rely on other EEG correlates such as the N1, which require knowledge of the target object’s temporal structure. Furthermore, hearing impaired individuals often have degraded object representations beginning at the level of auditory periphery (Shinn-Cunningham and Best, 2008; Dai et al., 2018) which may degrade the ability to use spatial features for focusing attention. Future work should explore the degree to which alpha is lateralized in hearing impaired listeners performing a spatial attention task.

In this study, we aimed to determine if lateralization of parietal alpha power reflected the use of spatial features during auditory selective attention. Our results showed that given a spatial cue, alpha was initially lateralized to reflect the location of the to-be-ignored auditory stream. We measured whether this lateralization would persist over the course of an auditory stream if strong pitch cues differentiated target from distractor. Our results showed that when pitch cues were strong, alpha lateralization was weakened after the target began to play, reflecting the fact that pitch could also be used to help focus attention. These results show that even when spatial attention is used initially to focus attention on a target, maintenance of attention can be accomplished using non-spatial cues when other acoustic features differentiate target from distractor streams.

## Acknowledgments

L.M.B, S.B., and B.G.S.-C. designed experiments and analyzed data. L.M.B. and S.B. performed experiments. L.M.B. and B.G.S.-C. wrote the paper. This work was supported by NIH Award No. 1RO1DC013825. L.M.B. was supported by NIH Computational Neuroscience Training Grant No. 5T90DA032484-05. The authors declare no conflicts of interest, financial or otherwise.

## References

Banerjee, S., Snyder, A. C., Molholm, S., and Foxe, J. J. (2011). Oscillatory alpha-band mechanisms and the deployment of spatial attention to anticipated auditory and visual target locations: supramodal or sensory-specific control mechanisms? Journal of Neuroscience, 31(27):9923–9932.

Braga, R. M., Wilson, L. R., Sharp, D. J., Wise, R. J., and Leech, R. (2013). Separable networks for top-down attention to auditory non-spatial and visuospatial modalities. NeuroImage, 74:77–86.

Brainard, D. H. (1997). The psychophysics toolbox. Spatial Vision, 10(4):433–436.

Cherry, E. C. (1953). Some experiments on the recognition of speech, with one and with two ears. The Journal of the Acoustical Society of America, 25(5):975–979.

Choi, I., Rajaram, S., Varghese, L., and Shinn-Cunningham, B. (2013). Quantifying attentional modulation of auditory-evoked cortical responses from single-trial electroencephalography. Frontiers in Human Neuroscience, 7:115.

Choi, I., Wang, L., Bharadwaj, H., and Shinn-Cunningham, B. (2014). Individual differences in attentional modulation of cortical responses correlate with selective attention performance. Hearing Research, 314:10–19.

Corbetta, M. and Shulman, G. L. (2002). Control of goal-directed and stimulus-driven attention in the brain. Nature reviews neuroscience, 3(3):201–215.

Dai, L., Best, V., and Shinn-Cunningham, B. G. (2018). Sensorineural hearing loss degrades behavioral and physiological measures of human spatial selective auditory attention. Proceedings of the National Academy of Sciences of the United States of America, 115(14):E3286–E3295.

Delorme, A. and Makeig, S. (2004). Eeglab: an open source toolbox for analysis of single-trial eeg dynamics including independent component analysis. Journal of neuroscience methods, 134(1):9–21.

Foxe, J. and Snyder, A. (2011). The role of alpha-band brain oscillations as a sensory suppression mechanism during selective attention. Frontiers in Psychology, 2:154.

Händel, B. F., Haarmeier, T., and Jensen, O. (2011). Alpha oscillations correlate with the successful inhibition of unattended stimuli. Journal of Cognitive Neuroscience, 23(9):2494–2502.

Hansen, J. C. and Hillyard, S. A. (1988). Temporal dynamics of human auditory selective attention. Psychophysiology, 25(3):316–329.

Kayser, J. and Tenke, C. E. (2006). Principal components analysis of laplacian waveforms as a generic method for identifying erp generator patterns: I. evaluation with auditory oddball tasks. Clinical Neurophysiology, 117(2):348–368.

Kayser, J. and Tenke, C. E. (2015). On the benefits of using surface laplacian (current source density) methodology in electrophysiology. International Journal of Psychophysiology, 97(3):171–173.

Kelly, S. P., Lalor, E. C., Reilly, R. B., and J., F. J. (2006). Increases in alpha oscillatory power reflect an active retinotopic mechanism for distracter suppression during sustained visuospatial attention. Journal of Neurophysiology, 95(6):3844–3851.

Kidd Jr, G., Favrot, S., Desloge, J. G., Streeter, T. M., and Mason, C. R. (2013). Design and preliminary testing of a visually guided hearing aid. The Journal of the Acoustical Society of America, 133(3):EL202–EL207.

Kong, Y.-Y., Mullangi, A., and Ding, N. (2014). Differential modulation of auditory responses to attended and unattended speech in different listening conditions. Hearing Research, 316:73–81.

Krumbholz, K., Nobis, E. A., Weatheritt, R. J., and Fink, G. R. (2009). Executive control of spatial attention shifts in the auditory compared to the visual modality. Human Brain Mapping, 30(5):1457–1469.

Larson, E. and Lee, A. K. (2014). Switching auditory attention using spatial and non-spatial features recruits different cortical networks. Neuroimage, 84:681–687.

Lee, A. K., Larson, E., Maddox, R. K., and Shinn-Cunningham, B. G. (2014). Using neuroimaging to understand the cortical mechanisms of auditory selective attention. Hearing Research, 307:111–120.

Lee, A. K. C., Rajaram, S., Xia, J., Bharadwaj, H., Larson, E., Hämäläinen, M. S., and Shinn-Cunningham, B. G. (2012). Auditory selective attention reveals preparatory activity in different cortical regions for selection based on source location and source pitch. Frontiers in Neuroscience, 6:190.

Lewis, J. W., Beauchamp, M. S., and DeYoe, E. A. (2000). A comparison of visual and auditory motion processing in human cerebral cortex. Cerebral Cortex, 10(9):873–888.

Maddox, R. K. and Shinn-Cunningham, B. G. (2012). Influence of task-relevant and task-irrelevant feature continuity on selective auditory attention. Journal of the Association for Research in Otolaryngology, 13(1):119–129.

Marrone, N., Mason, C. R., and Kidd, G. (2008). Evaluating the benefit of hearing aids in solving the cocktail party problem. Trends in amplification, 12(4):300–315.

McFarland, D. J. (2015). The advantages of the surface laplacian in brain–computer interface research. International Journal of Psychophysiology, 97(3):271–276.

Michalka, S. W., Kong, L., Rosen, M. L., Shinn-Cunningham, B. G., and Somers, D. C. (2015). Short-term memory for space and time flexibly recruit complementary sensory-biased frontal lobe attention networks. Neuron, 87(4):882–892.

Michalka, S. W., Rosen, M. L., Kong, L., Shinn-Cunningham, B. G., and Somers, D. C. (2016). Auditory spatial coding flexibly recruits anterior, but not posterior, visuotopic parietal cortex. Cerebral Cortex, 26(3):1302–1308.

Noyce, A. L., Cestero, N., Michalka, S. W., Shinn-Cunningham, B. G., and Somers, D. C. (2017). Sensory-biased and multiple-demand processing in human lateral frontal cortex. Journal of Neuroscience, 37(36):8755–8766.

Payne, L., Guillory, S., and Sekuler, R. (2013). Attention-modulated alpha-band oscillations protect against intrusion of irrelevant information. Journal of Cognitive Neuroscience, 25(9):1463–76.

Petersen, S. E. and Posner, M. I. (2012). The attention system of the human brain: 20 years after. Annual review of neuroscience, 35:73–89.

Posner, M. I. and Petersen, S. E. (1990). The attention system of the human brain. Annual review of neuroscience, 13(1):25–42.

Sauseng, P., Klimesch, W., Stadler, W., Schabus, M., Doppelmayr, M., Hanslmayr, S., Gruber, W. R., and Birbaumer, N. (2005). A shift of visual spatial attention is selectively associated with human eeg alpha activity. European Journal of Neuroscience, 22(11):2917–2926.

Shinn-Cunningham, B., Best, V., and Lee, A. K. (2017). Auditory object formation and selection. In The Auditory System at the Cocktail Party, pages 7–40. Springer.

Shinn-Cunningham, B. G. (2008). Object-based auditory and visual attention. Trends in cognitive sciences, 12(5):182–186.

Shinn-Cunningham, B. G. and Best, V. (2008). Selective attention in normal and impaired hearing. Trends in Amplification, 12(4):283–299.

Shomstein, S. and Yantis, S. (2006). Parietal cortex mediates voluntary control of spatial and nonspatial auditory attention. Journal of Neuroscience, 26(2):435–439.

Worden, M. S., Foxe, J. J., Wang, N., and Simpson, G. V. (2000). Anticipatory biasing of visuospatial attention indexed by retinotopically specific-band electroencephalography increases over occipital cortex. Journal of Neuroscience, 20(RC63):1–6.

